# Schwann cells have a limited window of time in which to initiate myelination program during early migration *in vivo*

**DOI:** 10.1101/2024.02.19.580938

**Authors:** Océane El-Hage, Aya Mikdache, Marie-José Boueid, Cindy Degerny, Marcel Tawk

## Abstract

The temporal control of mitotic exit of individual Schwann cells (SCs) is essential for radial sorting and peripheral myelination. However, it remains unknown when, during their multiple rounds of division, SCs initiate myelin signaling *in vivo*. By manipulating SC division during development, we report that when SCs skip their division during migration, but not during radial sorting, they fail to myelinate peripheral axons. This coincides with a sharp decrease in Laminin expression within the posterior lateral line nerve. Interestingly, elevating cAMP levels or forcing Laminin 2 expression within individual SCs restore their ability to myelinate, despite missing mitosis during migration. Our results demonstrate a limited time window during which migrating SCs initiate Laminin expression to activate the Laminin/Gpr126/cAMP signaling required for radial sorting and myelination at later stages *in vivo*.

## INTRODUCTION

Schwann cells (SCs) insulate axons with myelin sheaths to allow the rapid conduction of action potentials along myelinated nerves (Jessen and Mirsky, 2005). In order to associate with axons and become myelinating cells, SC precursors first migrate along the axons and divide while migrating before differentiating into immature SCs (Woodhoo and Sommer, 2008). Having reached their destination, SCs extensively proliferate and extend their processes to progressively radially sort large caliber axons (Boueid et al., 2020; Feltri et al., 2016; Mikdache et al., 2020). SCs then synthesize myelin proteins and wrap the large caliber axons with myelin sheath (Monk et al., 2015).

Several factors that regulate SC proliferation were identified and shown to play a major role during radial sorting. This includes cdc42, focal adhesion kinase (FAK), laminin 2 (Ln-2), laminin 8 (Ln-8), laminin γ1 and ErbB receptors (Benninger et al., 2007; Grove et al., 2007; Raphael et al., 2011; Yang et al., 2005; Yu et al., 2005). It is clear that the extracellular matrix protein Laminin is one major driver of SC development. Analysis of Laminin γ1-null SCs showed impaired proliferation and differentiation. Moreover, *Lama4^-/-^* and *Lama2^dy2J^* mutant mice showed a significant decrease in SC proliferation leading to amyelinating peripheral neuropathies. It has been proposed that Ln-2 and 8 promote the onset of radial sorting through proliferation. Another key signal in SC development is Neuregulin (Nrg1) which signals through ErbB receptors (Michailov et al., 2004). It regulates SC process extension during radial sorting in addition to its role in regulating SC numbers through proliferation (Raphael et al., 2011). Taken together, these results emphasize that the number of established SC at the onset of radial sorting is a key parameter regulating radial sorting, and consequently myelination.

Recently, we have shown that the temporal control of mitotic exit is essential for SC radial sorting and myelination (Mikdache et al., 2022). This process is regulated by the spindle pole protein Sil during both migration and radial sorting. However, it remains unknown whether divisions during both migration and radial sorting, are required for radial sorting and myelination and when individual SCs can initiate myelination signaling during their development *in vivo*. One possibility is that SCs migrate and divide along axons to reach their destination before going through extensive proliferation to regulate their numbers, deposit basal lamina (BL), radially sort axons and myelinate. Alternatively, SCs might initiate their myelination program while migrating and dividing by expressing and secreting their BL. The latter will contribute to myelination once SCs have finished migration.

Here, we used live imaging, pharmacological tools and manipulation of Laminin, Gpr126 and cAMP activities in zebrafish to specifically study SC divisions during migration and radial sorting and their impact on myelination *in vivo*. We show that individual SCs have a restricted window of time in which they initiate Laminin expression during migration, a key element used by SCs to activate the Gpr126/cAMP myelination pathway at later stages.

## RESULTS AND DISCUSSION

It is now well established that SCs first migrate and divide along growing axons of the posterior lateral line nerve (PLLn) between 24 and 48 hours post fertilization (hpf), and then intensively proliferate to radially sort and myelinate large caliber axons (Fig. 1A). Here, we use the larval zebrafish PLLn to monitor SC division during both migration and radial sorting. We show that all SCs go through several rounds of divisions, one during migration and at least one more during radial sorting. Of analyzed SCs, 58% (12/21) go through strictly two consecutive rounds of division up to myelination stages (72 hpf or 3 days post fertilization, dpf) (Fig. 1B-D’’, Table S1). The first division takes place during migration and the second during radial sorting. This data indicates that the majority of SCs experience mitosis during migration and radial sorting.

**Figure 1.**
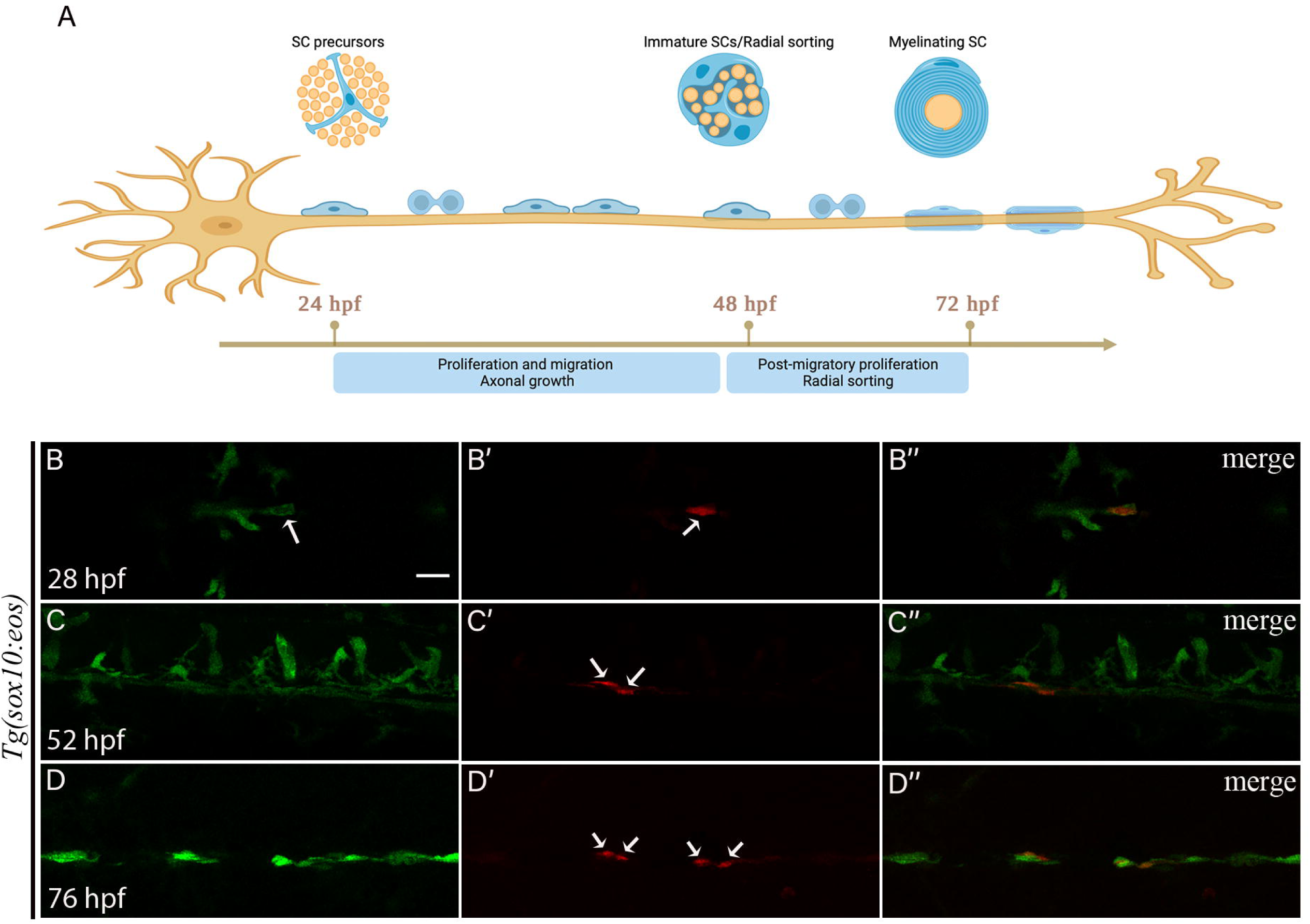
The majority of SCs divide during both migration and radial sorting. (A) SCs go through two main rounds of division: i) during migration and ii) during radial sorting. (B-B’’) a single SC after photoconversion (B, B’ white arrows). (C’) two red daughter cells are observed at 52 hpf (white arrows) and four cells at 76 hpf (white arrows in D’). B’’, C’’ and D’’: merge of B and B’, C and C’, D and D’ respectively. Scale bar = 20 μm.

In order to characterize the impact of SC division during migration and radial sorting on SC differentiation, we first carried out an ultrastructural analysis of PLLn in larvae treated with aphidicolin during radial sorting using transmitted electron microscopy (TEM). It is important to note that radial sorting and myelination start at the most anterior end of the PLLn and proceed gradually along its anterior-posterior (AP) axis. We therefore chose to study SCs around the region of the 7^th^ somite where radial sorting takes place after 48 hpf (Fig. S1). First, we incubated embryos in aphidicolin between 45 and 54 hpf (see Materials and Methods for aphidicolin treatment/efficacy and Fig. S2) to temporarily inhibit cell division during the initial phase of radial sorting. The medium was then washed and embryos were allowed to develop until 3 dpf. Another group of embryos was incubated from 45 hpf until 72 hpf to block SC division during the entire period of radial sorting. TEM analysis showed a significant decrease in the number of myelinated axons in both groups of treated embryos but also, as expected, in the total number of axons within the PLLn (Fig. 2A-E).

**Figure 2.**
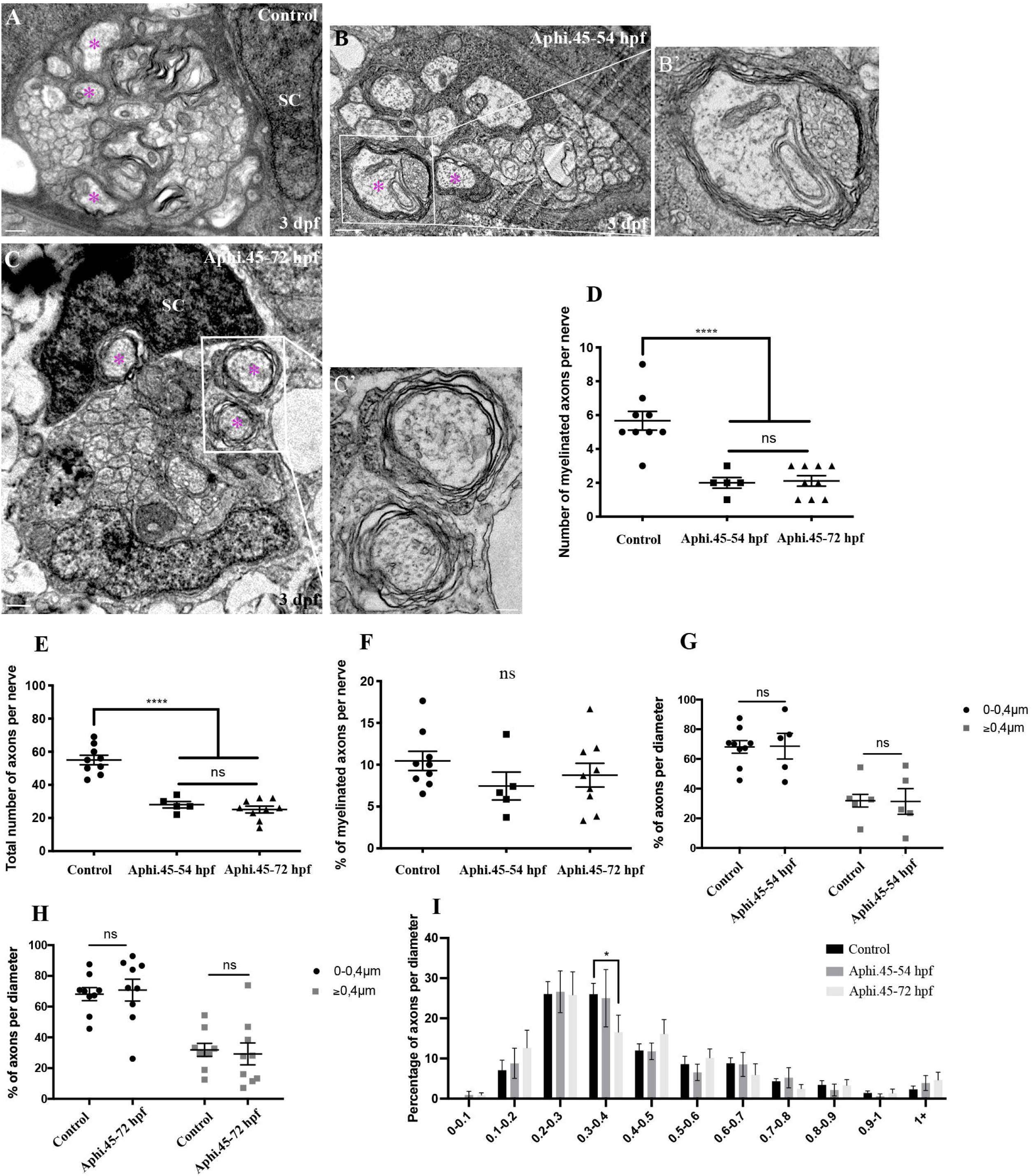
SC division during radial sorting is not required *per se* for myelination. TEM of a cross section of the PLLn in control (A), 45-54 hpf treated embryo (B) and 45-72 hpf treated embryo (C) at 3 dpf. Scale bars = 0.5 μm. Magenta asterisks represent some large caliber myelinated axons (some are shown at higher magnification in B’ and C’ (scale bars = 0.2 μm). (D) Quantification of the number of myelinated axons per nerve at 3 dpf in controls (average of 5.7±0.56, 9 nerves, n= 6 embryos), 45-54 hpf treated embryos (average of 2.00±0.31, 5 nerves, n= 5 embryos), and 45-72 hpf treated embryos (average of 2.11±0.32, 9 nerves, n= 5 embryos) (****, p≤0.0001; ns, p= 0.9790). (E) Quantification of the total number of axons per nerve at 3 dpf in controls (average of 57±2.9), 45-54 hpf treated embryos (average of 28.00±1.98) and 45-72 hpf treated embryos (average of 25.11±2.04) (****, p≤0.0001; ns, p= 0.6574). (F) Quantification of the percentage of myelinated axons relative to the total number of axons per nerve at 3 dpf in controls (average of 10.5±1.2), 45-54 hpf treated embryos (average of 7.46±1.66) and 45-72 hpf treated embryos (average of 8.76±1.42). ns, p> 0.05 in all cases. (G) Quantification of the percentage of axons according to their diameter relative to the total number of axons per nerve at 3 dpf in controls (average of 68.12 for 0-0.4 μm; 31.88 for >0.4 μm) and 45-54 hpf treated embryos (average of 68.62 for 0-0.4 μm; 31.38 for >0.4 μm). (H) Quantification of the percentage of axons according to their diameter relative to the total number of axons per nerve at 3 dpf in controls (average of 68.12 for 0-0.4 μm; 31.88 for >0.4 μm) and 45-72 hpf treated embryos (average of 70.76 for 0-0.4 μm; 29.24 for >0.4 μm). (I) Graph representing the distribution of axons relative to their diameter with 0.1 μm bin width at 3 dpf in controls, 45-54 hpf treated embryos and 45-72 hpf treated embryos (* p= 0.0234).

However, some large caliber axons were myelinated. Most importantly, the percentage of myelinated axons, the percentage of axons per diameter range and the distribution of axons relative to their diameter showed no significant difference between the different groups, except for a group of small caliber axons (Fig. 2F-I). This result suggests that cell division during radial sorting is dispensable for peripheral myelination.

Having previously identified an important role for *sil* in the temporal control of SC mitosis during migration, we sought to determine whether blocking cell division during migration might alter myelination. Other studies have established a role for ErbB receptors in SC division and migration and showed that cell division is not required for migration *per se* (Lyons et al., 2005). We therefore treated embryos with aphidicolin to block cell division specifically during SC migration (22 to 40 hpf; Movies S1, S2 and Fig. S3). The medium was then washed and embryos were allowed to develop until 3 dpf. As expected, there was a sharp decrease in the total number of axons per nerve, however, we counted an average of only 0.125 myelinated axon in treated embryos in comparison to 5.63 in control embryos (Fig. 3A-D). The percentage of myelinated axons per nerve showed a sharp decrease while the percentage of axons per diameter range and their distribution relative to their diameter were comparable to controls (Fig. 3E-G). The myelination defect correlated with a significant decrease in the number and percentage of radially sorted axons (Fig. S4). To test whether SCs were able to divide again once the medium was washed, we analyzed their numbers and division during radial sorting. We observed a significant decrease in the number of SCs in treated embryos but the number of PH3+ ones and the percentage of PH3+ SCs relative to the total number of SCs were similar to controls (Fig. S5A-E). We also observed a normal pattern of division during radial sorting in treated embryos similar to controls (Movies S3, S4 and Fig. S4F,G). We further analyzed treated embryos at later stages. We observed a sharp decrease in the number and percentage of myelinated axons per nerve at 5 dpf similar to what we observed at 3 dpf (Fig. S6A-E). Together, these data suggest that blocking cell division specifically during SC migration prevents myelination.

**Figure 3.**
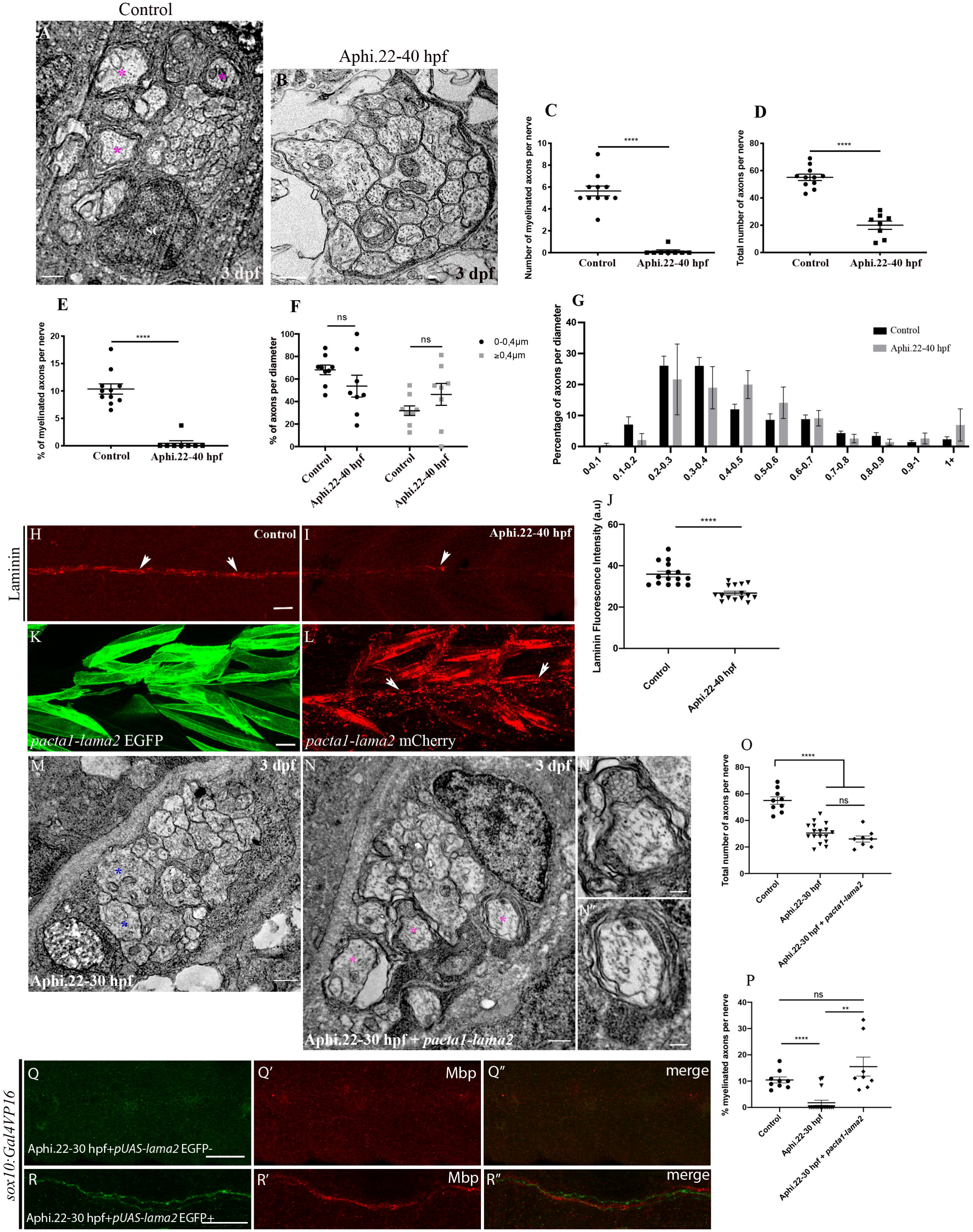
Blocking cell division during migration inhibits peripheral myelination and forcing Laminin expression restores myelination. TEM of a cross section of the PLLn in control (A) and 22-40 hpf treated embryo (B) at 3 dpf. Magenta asterisks represent some large caliber myelinated axons. Scale bars = 0.5 μm. (C) Quantification of the number of myelinated axons per nerve at 3 dpf in controls (average of 5.63±0.45, 11 nerves, n= 7 embryos) and 22-40 hpf treated embryos (average of 0.125±0.125, 8 nerves, n= 6 embryos) (****, p≤0.0001). (D) Quantification of the total number of axons per nerve at 3 dpf in controls (average of 55.09±2.32) and 22-40 hpf treated embryos (average of 20±3.04) (****, p≤0.0001). (E) Quantification of the percentage of myelinated axons relative to the total number of axons per nerve at 3 dpf in controls (average of 10.36±0.93) and 22-40 hpf treated embryos (average of 0.46±0.46) (****, p≤0.0001). (F) Quantification of the percentage of axons according to their diameter relative to the total number of axons per nerve at 3 dpf in controls (average of 68.12 for 0-0.4 μm; 31.88 for >0.4 μm) and 22-40 hpf treated embryos (average of 53.69 for 0-0.4 μm; 46.31 for >0.4 μm). (G) Graph representing the distribution of axons relative to their diameter with 0.1 μm bin width at 3 dpf in controls and 22-40 hpf treated embryos. Laminin expression in a control embryo (H) and treated embryo (I) at 48 hpf showing the PLLn nerve (arrows). Scale bar = 20 μm. (J) Quantification of Laminin fluorescence intensity along the PLLn in control (average of 36±1.4, n=15) and treated embryos (average of 26.80±0.92, n=15) at 48 hpf (****, p≤0.0001), a.u, arbitrary unit. (K) Lateral view of EGFP expression in muscles surrounding the PLLn at 48 hpf following *pacta1-lama2* injection. Scale bar = 20 μm. (L) Lateral view of mCherry-tagged secreted Laminin at 48 hpf in muscles and within the PLLn (white arrows). TEM of a cross section of the PLLn in treated embryo (M) and treated embryo injected with *pacta1-lama2* (N). Magenta asterisks represent some large caliber myelinated axons, (some are shown at higher magnification in N’,N’’, scale bars = 0.2 μm). Blue asterisks represent some large caliber non-myelinated axons. Scale bars = 0.5 μm. (O) Quantification of the total number of axons per nerve at 3 dpf in controls (average of 55.56±2.7, 9 nerves, n= 6 embryos), treated embryos (average of 30.53±1.75, 17 nerves, n=13 embryos) and treated embryos + *pacta1-lama2* (average of 26.00±2.32, 8 nerves, n=9 embryos) (****, p≤0.0001; ns, p=0.1401). (P) Quantification of the percentage of myelinated axons relative to the total number of axons per nerve at 3 dpf in controls (average of 10.73±1.4), treated embryos (average of 1.77±0.96) and treated embryos + *pacta1-lama2* (average of 15.53±3.61) (****, p≤0.0001; **, p= 0.0063; ns, p= 0.2166). Lateral view showing the absence of EGFP expression in SCs of the PLLn (Q) that correlates with a significant decrease in Mbp expression at 3 dpf (Q’), n= 20. Scale bar = 50 μm. Q’’, merge of Q and Q’. Lateral view showing EGFP expression in SCs of the PLLn (R) that correlates with normal Mbp expression at 3 dpf (R’) following *pUAS-lama2* injection with positive clones of EGFP in *sox10:Gal4VP16* 22-30 hpf treated embryos, n=16. R’’, merge of R and R’. Scale bar = 20 μm.

Our previous findings revealed an essential role for the temporal control of SC division in promoting radial sorting and myelination *in vivo* via Laminin2. We reasoned that SCs might have a limited window of time during migration in which they gradually set up the molecular signaling required for subsequent radial sorting and myelination. First, we tested whether blocking cell division during migration would alter Laminin expression within the PLLn as observed in *sil* mutants. Embryos incubated in aphidicolin during SC migration showed a significant decrease in the expression of Laminin along the PLLn at 48 hpf (Fig. 3H-J). Second, we treated embryos with aphidicolin for different periods of time during migration and we identified a time window of around 8 h in which SCs initiate myelination signaling during migration (Fig. 3M). To test if this temporal delay in division that results in myelination defects is related to a defective Laminin function, we injected embryos with 20 pg of *lama2* overexpression construct and treated embryos with aphidicolin between 22 and 30 hpf. The group of *lama2* injected/treated embryos showed a normal percentage of myelinated axons and of PH3+ SCs (Fig. 3K-Q; Fig. S7). We have previously shown that SCs are a significant source of Laminin for the PLLn, we therefore decided to force Laminin α2 expression within SCs and assess myelination in aphidicolin treated embryos. We injected 20 pg of *pUAS-lama2-EGF-mCherry* construct into *tg(sox10:Gal4VP16)* embryos to allow a specific expression of Lama2 within SCs. Injected/treated embryos that showed EGFP expression in SCs expressed normal levels of Mbp while those injected but were negative for EGFP showed a significant decrease in Mbp expression (Fig. 3R-S’’).

Laminin is a major component of the BL synthesized by SCs, and can mediate the Gpr126/cAMP signaling during SC development (Petersen et al., 2015). We therefore tested whether Laminin signals through Gpr126/cAMP to restore myelination in the absence of mitosis during migration. Embryos treated with aphidicolin during migration showed a significant decrease in *mbp* expression along the PLLn while those treated but injected with *lama2* overexpression construct or treated with forskolin (FSK) showed normal levels of *mbp* (Fig. 4A-C,F). In embryos that were injected with *gpr126* morpholino (MO) and treated with aphidicolin, the injection of *lama2* overexpression construct failed to restore *mbp* expression (Fig. 4B-C,D-G). These results suggest that Laminin interacts with Gpr126/cAMP pathway to restore myelination.

**Figure 4.**
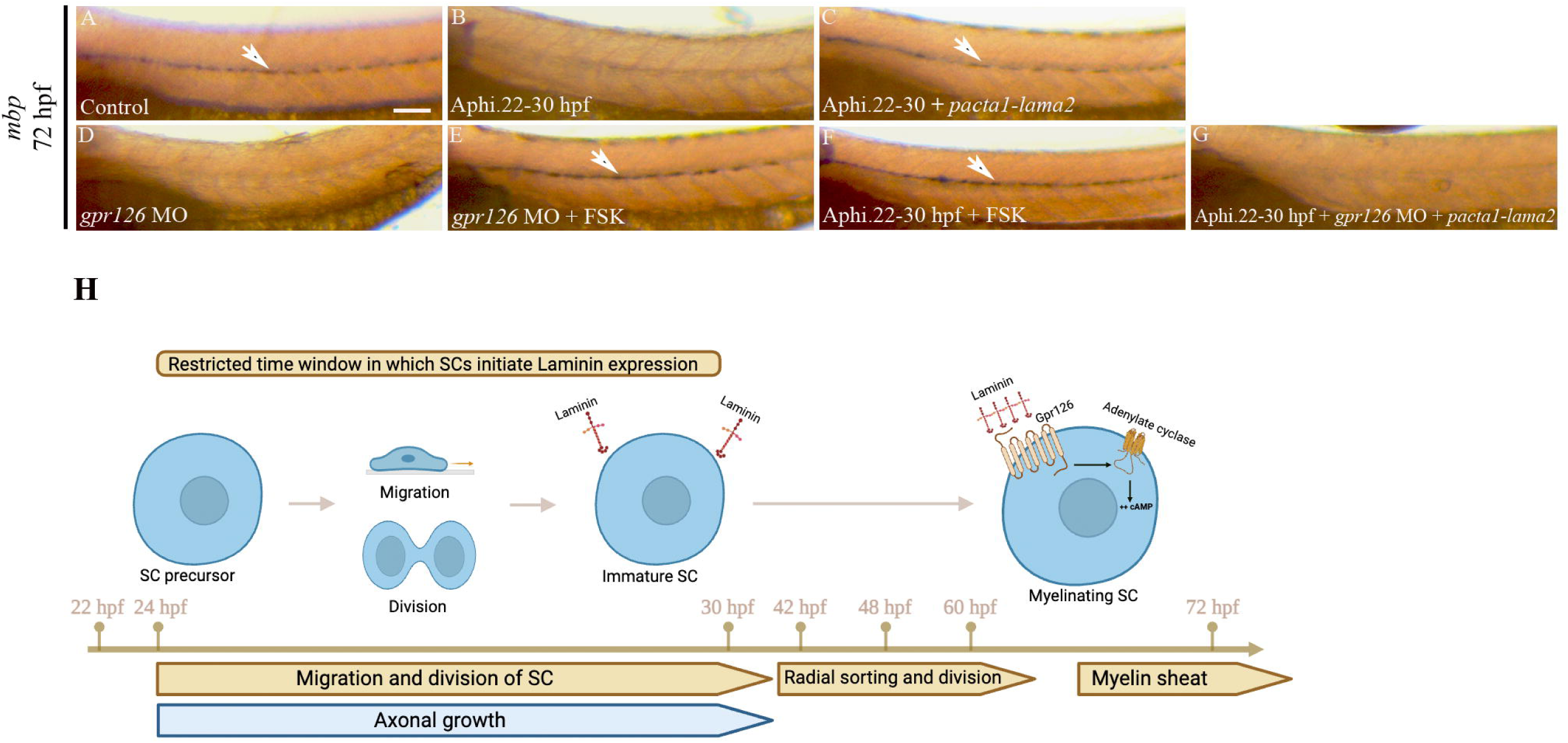
Laminin signals through Gpr126/cAMP to restore myelination in aphidicolin treated embryos. Lateral views of *mbp* expression at 3 dpf revealed by *in situ* hybridization along the PLLn in control showing a robust expression (A, n=32/32), aphidicolin treated embryos showing a sharp decrease in *mbp* expression (B, n=30/34), aphidicolin treated embryo and injected with *pacta-lama2* (C, n=37/38) showing a normal *mbp* expression in clones of SCs. *gpr126* morphants show a significant reduction in *mbp* expression (D, n=30/32) while those treated with FSK show a normal expression of *mbp* (E, n=27/30) similar to embryos treated with aphidicolin and FSK (F, n=31/33). Embryos treated with aphidicolin and injected with *pacta1-lama2* fail to express *mbp* in the absence of Gpr126 (G, n=30/34). Arrows indicate SCs expressing *mbp* along the PLLn. Scale bar = 200 μm. (H) Model of SC development, data strongly suggest a model in which SCs initiate Laminin expression during migration. Increasing levels and polymerization of Laminin drive radial sorting and myelination through Gpr126/cAMP signaling at later stages.

The key observation from this work is that skipping cell division during migration leads to defective myelination that correlates with a significant decrease in Laminin expression at the onset of radial sorting. Forcing Laminin expression in the vicinity of the PLLn or directly within SCs is enough to restore myelination. Aphidicolin treatment blocks cells in G1/S phase of the cell cycle in which DNA is replicated (Baranovskiy et al., 2014), which leads to collisions between transcription and replication machineries, impeding the expression or producing incomplete transcripts (Duch et al., 2013; Lin and Pasero, 2012). Moreover, when genes are replicated, there is a transient reduction in transcription activity (Wang et al., 2021). It is therefore possible that Laminin expression is directly altered following S phase blockade in SCs during migration, strongly suggesting that its expression is initiated within a limited window of time during SC migration.

In summary, our data support an *in vivo* model in which SCs need to express and secrete enough Laminin during migration. Laminin polymerization would in turn signal through the Gpr126/cAMP pathway to drive myelination at later stages. When cell division is delayed or prevented during migration, SCs fail to express Laminin and myelinate peripheral axons (Fig. 4H). Indeed, mutations in genes with cell cycle function are involved in Charcot-Marie-Tooth (CMT) disease and peripheral myelination defects (Mikdache et al., 2022; Timmerman et al., 2014). Taken together, our work provides some novel insights into the molecular process associated with cell division regulation in SCs during development and the potential implication of its deregulation in peripheral neuropathies.

## MATERIALS AND METHODS

### Embryo care

AB strain of zebrafish (*Danio rerio*) was used in this study. Embryos were staged and cared for according to standard protocols (https://zfin.org/zf_info/zfbook/cont.html). *Tg(foxd3:gfp)* stable transgenic line was used to label SCs (Gilmour et al., 2002). *Tg(sox10:Gal4VP16)^el159^*and *Tg(sox10:eos)^w9^* were kindly provided by Gage Crump (Center for stem cell and regenerative medicine, Keck School of medicine, University of South California, Los Angeles, CA, USA) and Sarah Kucenas (University of Virginia, Department of Biology, Charlottesville, VA, USA) respectively. All animal experiments were conducted with approved protocols at Inserm by DDPP Val de Marne, France under license number F 94-043-013.

### Microinjections

*pacta1-lama2* (kindly provided by Kelly Monk and Peter Currie) was injected at 20 pg per embryo along with 100 pg of Tol2 transposase mRNA. Embryos that showed EGFP expression in the muscles as well as mCherry expression within the vicinity of the PLLn were selected for further analysis. *pUAS-lama2* was injected into *Tg(sox10:Gal4VP16)^el159^* at 20 pg per embryo.

Embryos were injected at one cell stage with 15 ng of *gpr126* morpholino (5’-ACAGAATATGAATACCTGATACTCC-3’).

### In situ hybridization

*In situ* hybridization was performed following standard protocols using *mbp* probe. Embryos were raised in egg water with PTU (0.003%) to avoid pigmentation and were fixed overnight in 4% paraformaldehyde at 3 dpf for controls and following different treatments with aphidicolin. Embryos were dehydrated the next day and kept in methanol at −20°C. Embryos were then rehydrated and treated with proteinase K and incubated overnight with the *mbp* probe. Embryos were washed and the staining was revealed using anti-digoxigenin antibody.

### Immunofluorescence

The following antibodies and dilutions were used: rabbit anti-laminin (Sigma, 1/200; L9393), rabbit anti-phospho Histone 3 (Ser 10) (Millipore; 1/500; Cat # 06-570), rabbit anti-myelin basic protein (Custom produced (Tingaud-Sequeira et al., 2017), gift from Patrick Babin, 1/500), mouse anti-EGFP (Millipore; 1/500; REF MAB3580). Primary antibodies were detected with appropriate secondary antibodies conjugated to either Alexa 488, Alexa 568 at a 1:500 dilution. For immunostaining, embryos were fixed in 4% paraformaldehyde 1X PBS overnight at 4°C, then washed with 1xPBS to be dehydrated in methanol for at least 6 hours at −20°C. Samples were then rehydrated, digested with proteinase K and blocked with 0.5% triton in PBS and 10% sheep serum and then incubated with primary antibody overnight at 4°C (diluted in PBS+2% sheep serum). Larvae were then washed with PBS for a few hours and then incubated with secondary antibody in 0.5% triton in PBS and 2% sheep serum overnight at 4°C. Stained larvae were then imaged with a Leica SP8 confocal microscope. For Laminin and Mbp staining, embryos were fixed in 4% PFA for 30 min and treated for 7 min with acetone at −20°C. Samples were then blocked with 0.5% triton in PBS and 10% sheep serum and then incubated with primary antibody overnight at 4°C (diluted in PBS+2% sheep serum). Larvae were then washed with PBS for a few hours and then incubated with secondary antibody in 0.5% triton in PBS and 2% sheep serum for 3 hours at RT. For Laminin fluorescence intensity quantification, the same parameters (Excitation/emission, gain for detectors, lasers intensity) were applied for image acquisition in controls and aphidicolin treated embryos and fluorescence intensity was measured using image J (Analyze/measure).

### Photoconversion of SCs in *Tg(sox10:eos)^w9^*

SCs in *sox10:eos* transgenics were photoconverted in order to follow their development during migration and radial sorting. Embryos were first mounted at 28 hpf and imaged to detect any non-specific photoconversion. A region of interest (RO) that corresponds to a single SC was then exposed to UV light (1 minute) using the FRAP set up on Leica SP8 confocal. Embryos were imaged afterwards to confirm that single cells were photoconverted. Photoconverted SCs were then followed during two days and imaged every 24h after photoconversion.

### Aphidicolin and Forskolin treatment

Embryos were incubated in fish water containing 150 μM aphidicolin diluted in DMSO between 45 and 54 hpf or 45 and 72 hpf and with 100 μM between 22 and 40 hpf or 22 and 30 hpf. Controls were incubated in equivalent amount of DMSO (0.1%) solution during the same periods. It took around 2 to 3 h for the aphidicolin to become fully efficient following incubation.

Embryos treated with aphidicolin were very sensitive to forskolin (FSK), we therefore tested different times of incubation and were able to restore *mbp* expression in *gpr126* morphants following 3 h incubation in forskolin. In all experiments that required forskolin treatment, embryos were incubated in fish water containing 20 μM forskolin diluted in DMSO between 48 and 51 hpf. Controls were incubated in equivalent amount of DMSO solution (0.1%) during the same period.

### Acridine orange staining

Embryos were anesthetized with 0.03% tricaine and incubated in fish water containing 5 μM acridine orange for 20 minutes in the dark. After wash, they were embedded in 1.5% low melting point agarose, and imaged with a Leica SP8 confocal microscope. Transmitted light imaging was used to identify the PLLn and spinal cord and counting apoptotic corpses within these structures. Fluorescent corpses were counted at the level of the yolk extension.

### Schwann cells and PH3 positive SCs counts

*Tg(foxd3:gfp)* embryos as well as treated and injected ones were analyzed. SCs and PH3 positive SCs were counted from the most anterior to the end of yolk extension along the AP axis (≈900 μm) after immunostaining. SCs and PH3 positive SCs were counted blindly by two independent researchers using image J.

### Live imaging

Embryos were anesthetized with 0.03% tricaine and embedded in 1.5% low melting point agarose. Recordings were performed at 27°C using a Leica SP8 confocal microscope. Control *Tg(foxd3:gfp)* and treated *Tg(foxd3:gfp)* embryos were imaged at 28 hpf and 52 hpf and recordings were acquired for up to 8h.

### Transmission electron microscopy

3 and 5 dpf embryos were fixed in a solution of 2% glutaraldehyde, 2% paraformaldehyde and 0.1M sodium cacodylate pH 7.3 overnight at 4°C. This was followed by a post-fixation step in cacodylate-buffered 1% osmium tetroxide (OsO_4_, Serva) for 1h at 4°C and in 2% uranyl acetate for 1h at room temperature. The tissue was then dehydrated and embedded in epoxy resin. Sections were contrasted with saturated uranyl acetate solution and were examined with a 1010 electron microscope (JEOL) and a digital camera (Gatan).

### Statistical analysis

Means and standard deviations were calculated with Graph Pad Prism 7. All data were first tested for normal distribution using D’Agostino & Pearson normality test. All experiments with only two groups and one dependent variable were compared using an unpaired *t*-test with Welch’s correction if they passed normality test; if not, groups were compared with the nonparametric Mann-Whitney test. Statistically significant differences were determined using one-way ANOVA for all experiments with more than two groups but only one dependent variable. Error bars depict standard errors of the mean (SEM). ns, p>0.05; *, p≤0.05; **, p≤0.01; ***, p≤0.001; ****, p≤0.0001. n represents the number of embryos.

## Acknowledgments

We would like to thank Sarah Kucenas for *Tg(sox10:eos)^w9^* embryos and Jon Clarke for critical reading of the manuscript. This work was funded by Inserm and University Paris-Saclay.

## Supplementary information

**Figure S1. Radial sorting along the anterior-posterior (AP) axis of the PLLn**

(A) Lateral view of a 48 hpf embryo. The dotted line represents the AP position of the cross-section analysis by TEM. Anterior is to the left and dorsal to the top. Scale bar = 50 μm. (B) TEM of a cross-section of the PLLn of a WT embryo at 48 hpf. The PLLn is delineated in white dotted lines. Only 2 axons out of 94 (≈2%) were radially sorted at 48 hpf at this AP position (5 nerves, n=6 embryos). Scale bar = 0.5 μm.

**Figure S2. Analysis of PH3+ and acridine orange (AO)+ cells in aphidicolin treated embryos during radial sorting**

PH3 immunolabeling in *Tg(foxd3:gfp)* (A-A’’) and *Tg(foxd3:gfp)* embryos treated with aphidicolin between 45 and 48 hpf and analyzed at 48 hpf (B-B’’). Arrow in A’’ indicates a SC that is GFP and PH3 positive. A’’, merge of A and A’; B’’, merge of B and B’. Scale bar = 20 μm.

(C) Quantification of the number of PH3+ cells within a defined region of the PLLn at 48 hpf in control (average of 1.1±0.31, n=10) and aphidicolin treated embryos between 45 and 48 hpf (average of 0, n=7) (** p= 0.0067).

Acridine orange staining at 72 hpf in control (D-D’’) and aphidicolin treated embryos between 45 and 54 hpf (E-E’’) within a defined region of the PLLn. Scale bar = 20 μm. Arrowheads designate the PLLn in D’ and E’. D’’, merge of D and D’; E’’, merge of E and E’.

(F) Quantification of the number of AO positive cells in control (average of 0.25±0.13, n=12) and embryos treated with aphidicolin between 45 and 54 hpf (average of 0.33±0.19, n=12) within a defined region of the PLLn (ns, p= 0.67).

AO staining at 72 hpf in control (G-G’’) and aphidicolin treated embryos between 45 and 54 hpf (H-H’’) within a defined region of the spinal cord. G’’, merge of G and G’; H’’, merge of H and H’. Scale bar = 20 μm.

(I) Quantification of the number of AO positive cells in control (average of 0.66±0.31, n=12) and aphidicolin treated embryos between 45 and 54 hpf (average of 20.17±1.07, n=12) within a defined region of the spinal cord (****, p<0.0001). TL, transmitted light.

**Figure S3. Analysis of AO positive cells in aphidicolin treated embryos during migration**

Acridine orange staining at 52 hpf in control (A-A’’) and aphidicolin treated embryos between 22 and 40 hpf (B-B’’) within a defined region of the PLLn. Scale bar = 20 μm. Arrowheads designate the PLLn in A’ and B’. A’’, merge of A and A’; B’’, merge of B and B’.

(C) Quantification of the number of AO positive cells in control (average of 0.58±0.19, n=12) and embryos treated with aphidicolin between 22 and 40 hpf (average of 0.59±0.22, n=12) within a defined region of the PLLn (ns, p>0.9999).

AO staining at 52 hpf in control (D-D’’) and aphidicolin treated embryos between 22 and 40 hpf (E-E’’) within a defined region of the spinal cord. D’’, merge of D and D’; E’’, merge of E and E’. Scale bar = 20 μm.

(F) Quantification of the number of AO positive cells in control (average of 1.83±0.38, n=12) and aphidicolin treated embryos between 22 and 40 hpf (average of 48.42±1.74, n=12) within a defined region of the spinal cord (****, p<0.0001). TL, transmitted light.

**Figure S4. Analysis of radial sorting in aphidicolin treated embryos between 22 and 40 hpf**

(A) Quantification of the number of sorted axons per nerve at 3 dpf in control (average of 7±0.5) and aphidicolin treated embryos between 22 and 40 hpf (average of 0.5±0.3) embryos (**** p≤0.0001).

(B) Quantification of the percentage of sorted axons per nerve at 3 dpf in control (average of 12.97±1.21) and aphidicolin treated embryos between 22 and 40 hpf (average of 2.8±2.3) (**p= 0.0027).

**Figure S5. Analysis of SC division during radial sorting in embryos treated with aphidicolin between 22 and 40 hpf**

PH3 immunolabeling at 52 hpf in *Tg(foxd3:gfp)* (A-A’’) and *Tg(foxd3:gfp)* embryos treated with aphidicolin between 22 and 40 hpf (B-B’’). Arrow in A’’ and B’’ indicate SCs that are GFP and PH3 positive. A’’, merge of A and A’; B’’, merge of B and B’. Scale bar = 20 μm.

(C) Quantification of the number of Schwann cells within a defined region of the PLLn at 52 hpf in control (average of 59.10±2.44 cells, n= 10 embryos) and embryos treated with aphidicolin between 22 and 40 hpf (average of 18±3.27 cells, n= 11 embryos) (****, p≤ 0.0001). (D) Quantification of the number of PH3^+^/Schwann cells within a defined region of the PLLn at 52 hpf in control (average of 0.87±0.47, n=8 embryos) and embryos treated with aphidicolin between 22 and 40 hpf (average of 0.18±0.12, n= 11 embryos) (ns, p= 0.3265).

(E) Quantification of the percentage of PH3^+^ Schwann cells relative to the total number of Schwann cells within a defined region of the PLLn at 52 hpf in control (average of 2.54±0.95, n= 10 embryos) and embryos treated with aphidicolin between 22 and 40 hpf (average of 2.34±1.65, n= 11 embryos) (ns, p= 0.3274).

(F) Still images of time-lapse imaging at 52 hpf in *Tg(foxd3:gfp)* and *Tg(foxd3:gfp)* treated with aphidicolin between 22 and 40 hpf embryos. Arrows indicate Schwann cells along the PLLn at different timepoints prior to and after division. Scale bars = 20 μm.

(G) Quantification of the time required for control (average of 8.70±0.41 min, 20 cells, n= 5 embryos) and aphidicolin treated embryos between 22 and 40 hpf (average of 8.14±0.67 min, 7 cells, n= 4 embryos) Schwann cells to successfully complete mitotic division during radial sorting. (ns, p= 0.1796).

**Figure S6. Analysis of the ultrastructure of the PLLn at 5 dpf in aphidicolin treated embryos between 22 and 30 hpf**

TEM of a cross section of the PLLn at 5 dpf in control (A) and embryo treated with aphidicolin between 22 and 30 hpf (B). Scale bars = 0.5 μm. Magenta asterisks represent some large caliber myelinated axons. Some large caliber axons are shown at higher magnification in A’ and B’. Scale bars = 0.2 μm.

(C) Quantification of the number of myelinated axons per nerve at 5 dpf in controls (average of 9±1.29, 4 nerves, n= 4 embryos) and embryos treated with aphidicolin between 22 and 30 hpf (average of 0, 4 nerves, n= 4 embryos) (**p= 0.0061).

(D) Quantification of the total number of axons per nerve at 5 dpf in controls (average of 49.75±3.32) and embryos treated with aphidicolin between 22 and 30 hpf (average of 28.75±3.27) (**p=0.0041).

(E) Quantification of the percentage of myelinated axons relative to the total number of axons per nerve at 5 dpf in controls (average of 17.94±1.92) and embryos treated with aphidicolin between 22 and 30 hpf (average of 0) (**p= 0.0021).

**Figure S7. Analysis of the number of SCs and their proliferation in aphidicolin treated embryos between 22 and 30 hpf following *pacta1-lama2* injection**

(A) Quantification of the number of SCs within a defined region of the PLLn at 48 hpf in controls (average of 72.8±2.71, n= 10 embryos), controls injected with *pacta1-lama2* (average of 48.8±2.84, n= 10 embryos), aphidicolin treated embryos between 22 and 30 hpf (average of 19.44±1.10, n= 9 embryos) and aphidicolin treated embryos between 22 and 30 hpf injected with *pacta1-lama2* (average of 21±1.49, n= 10 embryos). (ns p>0.9999).

(B) Quantification of the number of PH3+ SCs within a defined region of the PLLn at 48 hpf in controls (average of 1±0.26, n= 8 embryos), controls injected with *pacta1-lama2* (average of 1±0.25, n= 10 embryos), aphidicolin treated embryos between 22 and 30 hpf (average of 0.44±0.17, n= 9 embryos) and aphidicolin treated embryos between 22 and 30 hpf and injected with *pacta1-lama2* (average of 0.7±0.21, n= 10 embryos).

(C) Quantification of the percentage of PH3+ SCs relative to the total number of SCs within a defined region of the PLLn at 48 hpf in controls (average of 1.35±0.38, n= 8 embryos), controls injected with *pacta1-lama2* (average of 2.07±0.54, n= 10 embryos), aphidicolin treated embryos between 22 and 30 hpf (average of 2.13±0.86, n= 9 embryos) and aphidicolin treated embryos between 22 and 30 hpf and injected with *pacta1-lama2* (average of 4±1.58, n= 10 embryos).

**Movie S1. Real-time imaging of SCs in *Tg(foxd3:gfp)* embryo at 28 hpf**

A 28 hpf embryo expressing GFP in SCs, the control embryo was imaged every 5 minutes for several hours by confocal microscopy. Lateral view, anterior to the left and dorsal to the top. Arrows indicate SCs in migration and arrowheads indicate SCs before and after division. This video represents five hours of continuous real-time imaging.

**Movie S2. Real-time imaging of SCs in *Tg(foxd3:gfp)* embryo at 28 hpf treated with aphidicolin at 22 hpf**

A 28 hpf embryo expressing GFP in SCs, the aphidicolin treated embryo was imaged every 5 minutes for several hours by confocal microscopy. Lateral view, anterior to the left and dorsal to the top. Arrows indicate SCs in migration and arrowheads indicate two adjacent SCs, note the distance that separate these two (arrowheads). This video represents five hours of continuous real-time imaging.

**Movie S3. Real-time imaging of SCs in *Tg(foxd3:gfp)* embryo at 52 hpf**

A 52 hpf embryo expressing GFP in SCs, the control embryo was imaged every 3 minutes for several hours by confocal microscopy. Lateral view, anterior to the left and dorsal to the top. Arrows indicate SCs before and after division. This video represents three hours and 20 minutes of continuous real-time imaging.

**Movie S4. Real-time imaging of SCs in *Tg(foxd3:gfp)* embryo at 52 hpf treated with aphidicolin between 22 and 30 hpf**

A 52 hpf embryo expressing GFP in SCs, the aphidicolin treated embryo was imaged every 3.5 minutes for several hours by confocal microscopy. Lateral view, anterior to the left and dorsal to the top. Arrows indicate SCs before and after division. This video represents four hours of continuous real-time imaging.

**Table S1. Analysis of SC development following photoconversion in *Tg(sox10:eos)^w9^* embryos**

Yellow lines depict SCs that went through strictly two rounds of division during their development (up to 76 hpf), one during migration and another during radial sorting. * 1 division per cell; ^#^ more than 1 division per cell.

